# Can Random Walking on a Hi-C Contact Matrix Lead to Data Quality Improvement? An Assessment

**DOI:** 10.1101/2025.06.11.659235

**Authors:** Yongqi Liu, Shili Lin

**Affiliations:** Department of Statistics, The Ohio State University, Columbus, OH 43210

**Keywords:** Hi-C, smoothing, random walk, random walk with restart, Topologically associated domains

## Abstract

Hi-C and single cell Hi-C (scHi-C) data are now routinely generated for studying an array of biological questions of interest, including whole genome chromatin organization to gain a better understanding of the chromosome three-dimensional hierarchical structure: compartments, Topologically Associated Domains (TADs), and long-range interactions. Due to concerns about data quality, especially for scHi-C because of its sparsity, data quality improvement is seen as a necessary step before performing analyses to answer biological questions. As such, methods have been developed accordingly, among them is a set of methods that are “random walk”-based, including random walk with a limited number of steps (RWS) and random walk with restart (RWR). Nevertheless, there is little justification for the use of such methods, nor quantification of their performance success. Taking correct identification of TADs as the end point, in this paper, we describe the characteristics of random-walk-based approaches and carry out empirical investigation for identifying TADs before and after random walks. Due to the lack of practical guidelines for choosing tuning parameters necessary for performing random walks, it is difficult to know how many steps of random walk for RWS or how small a restart probability for RWR should one choose to achieve good performance. Even in the unrealistic scenario when one has the hindsight of using the optimal parameter values, little improvement in downstream studies by first performing random walk was observed. This conclusion was based on extensive analytical analyses, simulation study, and real data applications. Therefore, the current study provides a cautionary note to researchers who may consider using random-walk-based approaches prior to downstream analyses.

## 1 Introduction

Chromosome conformation capture (3C) and its derivatives are methods for quantifying interactions among genomic loci. In particular, the High-throughput Chromosome Conformation Capture (Hi-C) technology [1] and other subsequent improvements [2, 3] permit genome-wide studies of the chromatin three-dimensional (3D) structure. Numerous methods have been developed in the past decade to process Hi-C data (bulk or single-cell) with various goals. One prominent example is for finding Topologically Associated Domains (TADs), which are contiguous regions of the genome associated with chromatin 3D structures and functions [4]. Loci within the same TAD interact with much higher frequency than those between different TADs, and the domain boundaries are often demarcated by CTCF insulator proteins [5, 6].

With the development of the technology, single-cell Hi-C data (scHi-C) are available for more in-depth studies of chromatins [7–11]. Hundreds or even thousands of cells can now be processed in a single run, which can then be used to study cell-to-cell variability in chromatin structures for cells within the same cell type. For example, Nagano et al. (2017) [10] utilized thousands of scHi-C data to study the chromosome cell-cycle dynamics and place the cells along the cell-cycle.

Hi-C data are usually organized as an *N* × *N* contact matrix (*M*), where *N* is the number of equal-sized bins of the genome (also referred to as loci). Each element of the contact matrix is the number of pair-end reads signifying interactions between the corresponding pair of loci. Sparsity (i.e. with many zeros in the contact matrix) is a major concern, especially for scHi-C data [12, 13], where the sequencing depth is often less than one-tenth of bulk data [11]. For better performance of downstream analysis, such as revealing the underlying domain structures, data quality improvement is essential. To address this issue, a number of methods have been developed, which all aim to smooth the data in some way and impute the zeros by borrowing information from neighbors. Such methods include HiCRep [14], HiCPlus [15], GenomeDISCO [16], scHiCluster [12], SCL [17], DeepHiC [18], ScHiC-Rep [19], SnapHiC [20], SnapHiC2 [21], scHiCStackL [22], Higashi [23], HiCImpute [24], and scHiCPTR [25]. Some of these methods were proposed for bulk while some were specifically for single cell data where further information may be borrowed from similar single cells and even bulk data. Among them, a prominent class is random-walk based [16, 12, 19–22, 25]. Supplementary Table S1 provides a list of methods/software packages that aim to address the issue of sparsity and data quality improvement, where the first half contains methods that are (at least partially) random-walk based.

A standard random walk method with a finite number (say *s*) of steps (RWS) propagates the information of all interactions related to locus *i* and all interactions related to locus *j* through the transition matrix constructed from the contact count matrix. The (*i, j*)^th^ element of an RWS-smoothed contact matrix is the corresponding element in the *s*-step transition probability matrix (TPM), for a suitable *s*. It is suggested in GenomeDISCO that performing three steps of RWS (*s* = 3) gives the best result of smoothing with the purpose of quantifying reproducibility of data [16]. As for scHiCluster and SnapHiC, part of their data quality improvement methods is to conduct a random walk with restart (RWR) algorithm, which utilizes not only the global information as in RWS but also preserve some local information through repeated restart [12, 20].

The idea of using RWR for Hi-C data smoothing was borrowed from the usage of RWR in image processing, which appears to have emerged from the field of Computer Science [26]. In Pan et al. [26], the authors defined the affinity between two nodes in a network (analogous to the interaction intensity of two genomic loci in Hi-C data) to be the “steady-state” probability of RWR, which is the usual stationary distribution in a random walk modified by a restart component. In a standard random walk, the stationary distribution can be found by iteratively taking the product of the one-step transition matrix; similarly, the steady-state probability of RWR is found by taking many iterations until convergence. Therefore, unlike the use of RWS proposed in [16] for a finite number of steps, RWR generally runs the algorithm until convergence.

For methods relying on RWS, although random walking for three steps was suggested for one particular goal [16], whether this is appropriate for other purposes with different datasets has not been investigated. On the other hand, there does not seem to be any concrete recommendation for the restart probability in RWR, to the best of our knowledge. SnapHiC uses restart probability of 0.05 without justification [20], while scHiCluster claims in the appendix that this method is robust to the choice of restart probability, where the values of 0, 0.1, 0.2, …, 0.9 were tested [12]. Despite the prevalence of random-walk-based methods in Hi-C research, the rationale behind RWS and RWR (and the corresponding obligatory tuning parameter selection) for data quality improvement remains unclear, and little has been done to evaluate their impact on downstream analyses.

To apply a random-walk method, a Hi-C data matrix needs to be turned into a TPM. For Hi-C data manipulation, the first step is usually bias removal. This normalization can be done by vanilla coverage (VC), which divides a contact count by the sum of the corresponding row and column [1], square root vanilla coverage (sqrtVC), which divides a contact count by the square root of the product of corresponding row sum and column sum [2], or iterative correction and eigenvector decomposition (ICE), which iteratively corrects for the experimental biases and then performs eigenvalue decomposition [27]. Each element in the bias-removed matrix is then divided by the row sum, so that the sum of each row now equals to one, the characteristic of a TPM. However, such a TPM is no longer symmetrical, violating the basic property of a Hi-C derived data matrix.

In this study, we aim to investigate whether random-walk-based methods are good choices for improving Hi-C data quality, bulk or single cells. We first study the theoretical basis of RWS and RWR for Hi-C data quality improvement and carry out empirical study to visualize the data before and after random walking on the Hi-C matrix. Then, taking detecting TADs as an example downstream analysis, we investigate the impact of RWS- and RWR-smoothed data matrices on performing such a task. In particular, we consider three TADs detection algorithms from a large collection of such methods, including CaTCH [29], HiCseg [30], and TopDom [31], which were top performers among twenty-two TAD finders from a review article [32]. As a by-product, we also evaluate the relative performance of these three methods to assess the claims of robustness and consistency drawn in the review paper.

## 2 Materials and Methods

We first discuss the theory of RWS and RWR in the context of Hi-C contact matrices with TAD structures; that is, greater contacts among loci within a TAD (diagonal blocks in the contact matrix) but much weaker contacts between TADs. The properties of RWS can be easily found in textbooks on stochastic processes (e.g. [32]), as RWS is simply a special case of a Markov chain. On the other hand, the concept of RWR is relatively new and primarily used in computer science, and its property is not widely studied, to the best of our knowledge. Therefore, in this paper, we will study the stationary distribution of RWR within our context. Since a Hi-C matrix is symmetric, we implement the Knight-Ruiz (KR) algorithm [33] for normalization to obtain a doubly stochastic TPM that preserves such a symmetric property.

### 2.1 The theory of RWS as a special case of Markov chains

Suppose *P* is an *N* × *N* TPM. Then the (*i, j*)^th^ element of the transition matrix represents the probability of a RWS starting at locus *i* reaching locus *j* in one step. The *s*-step TPM *P*^(*s*)^ = *P*^*s*^ is obtained by multiplying *s* one-step TPM together. It is easily seen that *P*^(*s*)^ is also symmetrical if we start with a symmetrical TPM *P*; therefore, it is doubly stochastic. Suppose the limiting matrix *P*\* exists, then it can be obtained by letting *s* → ∞ (or setting *s* to be sufficiently large in practice) and the limiting matrix will have identical rows (or nearly identical rows for a finite but large enough *s*) — each row is referred to as the limiting distribution. Further, if the TPM *P* corresponds to an irreducible Markov chain (i.e. every locus can be reached from every other locus), then the limiting matrix will in fact have all identical entries. In the context of our problem, where our goal is to improve data quality — preserving TADs and enhancing their detections for example — we would not want to take a long random walk to reach the limiting matrix, since TADs would disappear. Therefore, the essential question is the goldilocks number, not too few but not too many steps. We will explore whether a rule of thumb provided in the literature, three steps (i.e. *s* = 3), works in various settings.

We provide a concrete example where *P* is a symmetric doubly stochastic matrix (can be thought of as a KR-normalized matrix from a Hi-C contact matrix — see Section 2.3) with TAD structures:

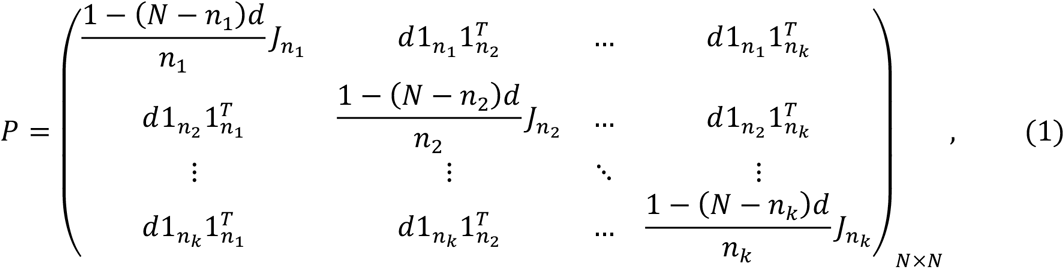

where we hypothesize *k* TADs each of size 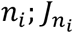is an all-1 square submatrix of dimension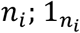is an all-1 column vector of length *n*_*i*_, *i* = 1, …, *k*, and *d* ≥ 0 is a constant. Thus, the total number of loci is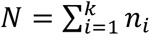. Since *P* is doubly stochastic, the limiting matrix is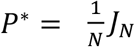 when *d* > 0. This simple example demonstrates that, if too many steps are taken using RWS, the smoothed matrix will simply merge all domains together and the original block-diagonal structure is ruined. The first row of the heatmaps in Supplementary Figure S1 show *P*^(*s*)^, for *s* = 1, 2, 3 and 10 for an ideal matrix as in (1): *N* = 200, *k* = 5, *n*_*i*_ = 50, 30, 20, 90, 10 for *i* = 1, …, 5, respectively, and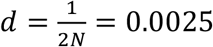. We can see that, after 10 steps, all domains vanished, illustrating the above theoretical analysis.

### 2.2 The limiting matrix of RWR for Hi-C matrix with TAD structures

Suppose we start with the same TPM *P* as in RWS. Then, for RWR, at each locus *i*, there is an additional fixed probability *α* ∈ [0,1] that the Markov chain can restart; that is, remaining at locus *i*. The *s*-step TPM *P*^(*s*)^ = (1− *α*)*P*^(*s*−1)^*P* + *αI*_*N*_, *s* = 1, 2, …, where we set *P*^(0)^ = *I*,, the identity matrix with the same dimension as the TPM *P*. We can easily see that *P*^(*s*)^ is symmetrical if *P* is symmetrical. Further, when *α* = 0, *P*^(*s*)^ = *P*^*s*^, reducing to RWS with *s* steps. Suppose the limiting matrix *P*\* exists, then it must satisfy *P*\* = (1 − *α*)*P*\**P* + *αI*_*N*_, which, after some simplification, can be expressed as

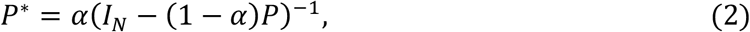

provided that the inverse exists.

Considering the same example matrix as defined in (1), we obtain the limiting matrix for RWR:

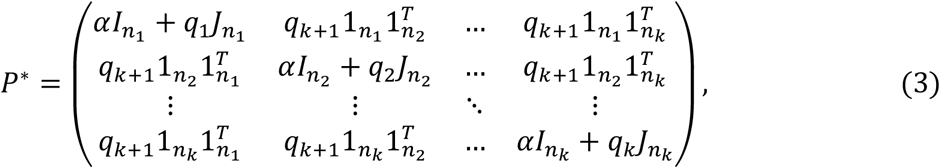

where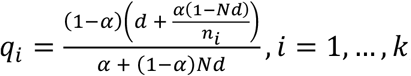, and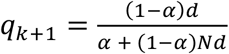. First, we note that, when *α* = 0, *P*\* is the same as the limiting distribution of a standard random walk when *d* > 0. In the other extreme, when *α* = 1, it can be seen that *P*\* = *I*_*N*,_ an identify matrix, thus also losing the domain structure. For a proper 0 < *α* < 1, the magnitude of *q*_*i*_, *i* = 1, …, *k* + 1, are all comparable, and roughly in the order of 1/*N*. On the other hand, although *P*\* is still written in a block-diagonal form, the elements in the main diagonal,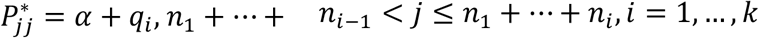(denoting *n*_0_ = 0), is dominated by *α* (*α* is typically in the order of one tenth whereas *q*_*i*_ is typically at least an order of magnitude lower for Hi-C data). As such, the most prominent feature of the limiting matrix is the main diagonal only, not the blocks. As *α* gets larger, the diagonal feature becomes more obvious, not surprisingly given the property when *α* = 1. The second row of the heatmaps in Supplementary Figure S1 shows *P*\* for values of *α* = 0.05, 0.1, 0.2, 0.5. We can see that, regardless of the *α* values, the most prominent feature of the limiting matrix is in fact the main diagonal, substantiating our theoretical analysis for RWR.

### 2.3 Implementation of KR normalization

As a preprocessing step, the Knight-Ruiz algorithm [33] is utilized for normalizing an input Hi-C data matrix before applying RWS and RWR. A typical Hi-C data matrix is symmetrical, with every element in the matrix being a non-negative integer. Therefore, the KR algorithm is particularly suited for transforming such a matrix into a doubly stochastic TPM that preserves these characteristics. Let *M* denote an *N* × *N* Hi-C count data matrix, where all the entries in the matrix are non-negative integers. The KR algorithm finds an *N*-dimensional diagonal matrix *D* = diag(*x*\*), where *x*\* is a vector of positive numbers such that matrix *P* = *DMD* is doubly stochastic. We note in passing that this matrix balancing problem can be written as solving a system of linear equation involving *x*\*, and an inner-outer iteration scheme was proposed [33] for solving the equations. The matrix *P* is symmetric with non-negative entries, and elements in each row (and each column) sums to one. In real data analysis, sometimes there are all-0 rows (and columns) in a contact matrix. In this case, we replace the diagonal elements in those rows with 1 before performing KR normalization. This manipulation, instead of cutting out all-zero rows (and columns), avoids creating fictitious TADs.

## 3 Results

### 3.1 Simulation Studies

We present three simulation studies to illustrate the behavior of the RWS- and RWR-smoothed matrices and their impact on downstream TAD detections. These studies range from an ideal Hi-C data matrix with contrived TAD structures, to a realistic simulation setting, to simply subsampling from a real Hi-C data matrix. In addition to heatmap visualization, we also use the Adjusted Rand Index (ARI) [34], a number between −1 and 1, as an objective measure of the congruence between the “true” TADs and the detected TADs, with a higher number denoting a greater congruence. This is done by assuming loci within each TAD in the “ground truth” structure belong to a cluster. The TADs identified from a data matrix are sorted into clusters the same way and compared to the “true clusters” for computing the ARI.

#### 3.1.1 Study 1: An idealized Hi-C data matrix with TADs

Our first study is based on the ideal TPM *P* in (1) with the same parameter values as in Supplementary Figure S1 except that we set *d* = 0. We simulated noise at each position of the matrix by drawing it randomly from the Unif(0, 0.02) distribution and added it to *P*, leading to the observed matrix *P*_*obs*_ = *P* + *R*. We then KR normalized it to obtain the doubly stochastic matrix *P*_*KR*_. We applied RWS to *P*_*KR*_ for *s* = 2, 3, 4, 5, and 10 steps, with the resulting smoothed matrices denoted as RWS2s, RWS3s, RWS4s, RWS5s, and RWS10s, respectively. For RWR, we considered several restart probabilities, *α* = 0.05, 0.1, 0.2, and 0.5, with the resulting smoothed matrices denoted as RWR0.05, RWR0.1, RWR0.2, and RWR0.5, respectively. A representative matrix from each of the two random-walk smoothing methods, together with the data matrices, are shown in the first column of Figure 1; all the smoothed matrices are in Supplementary Figures S2 and S3.

**Figure 1.**
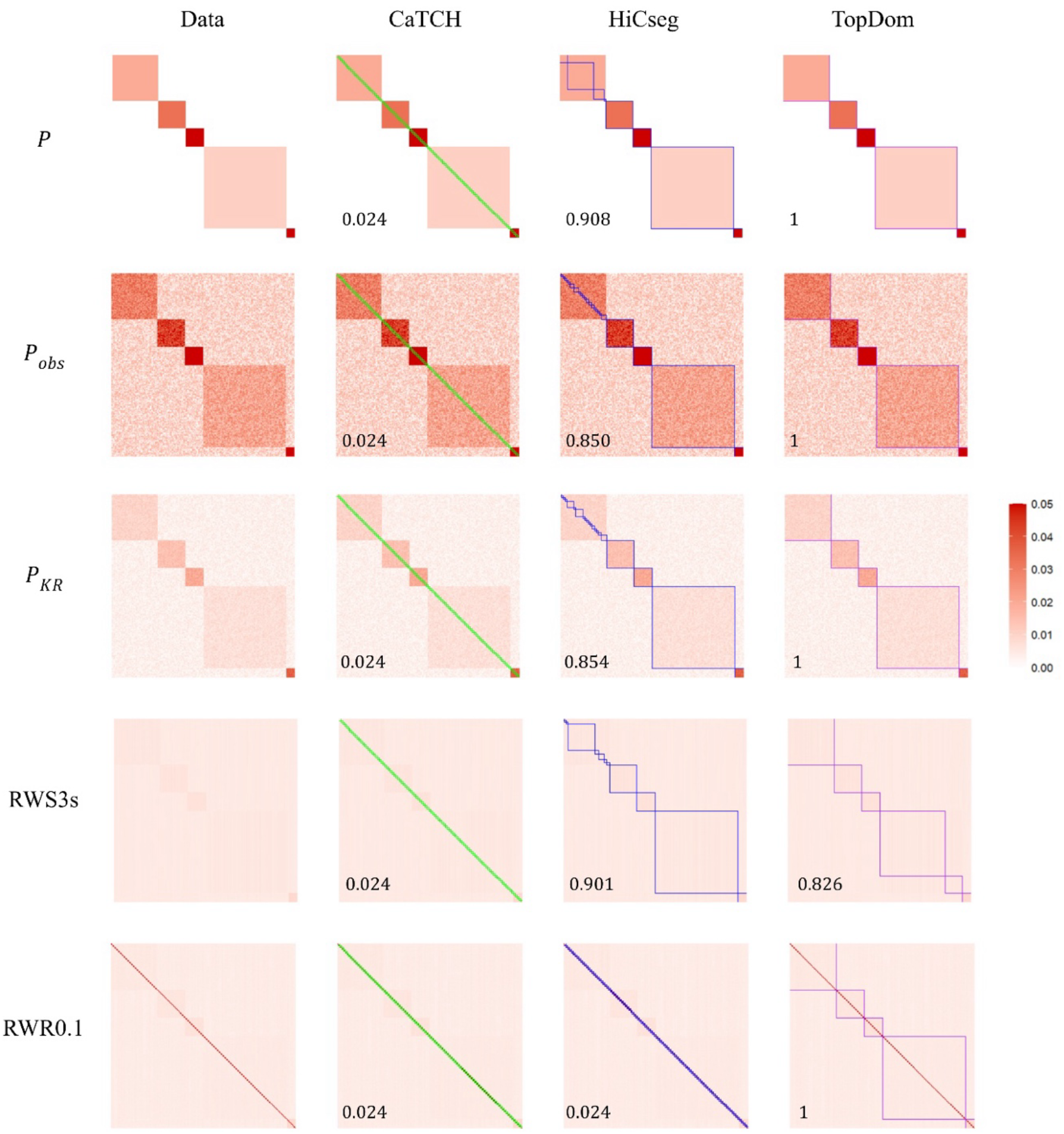
Heatmap visualization of the data matrices, along with the detected domain boundaries and ARI values (bottom left corner). The five rows give various kinds of data matrices: the perfect block diagonal data (*P*), the observed data with noise (*P*_*obs*_), the KR-normalized data matrix (*P*_*KR*_), the RWS-smoothed matrix with 3 steps (RWS3s), and the RWR-smoothed matrix with restart probability 0.1 (RWR0.1). The four columns show the data matrix (1^st^ column), the data matrix superimposed with the domain boundaries from CaTCH (2^nd^ column, green boxes), HiCseg (3^rd^ column, blue boxes), and TopDom (4^th^ column, purple boxes). The color scheme for all the heatmaps ranges from 0 (white) to 0.05 (red), with those values that are greater than 0.05 censored at 0.05.

We observed that the domain boundaries become a bit blurry but are still visible with the KR-normalized matrix. For RWS, other than the one with just two steps (RWS2s), the domains are difficult to discern or complete gone; the resulting smoothed matrix is simply uniform across all rows and columns. For RWR, with the entire range of restart probability, the most prominent feature is the diagonal line, which has much higher interaction intensities compared to its surrounding entries, although the some of the domain boundaries are still visible. These observations imply that applying RWS and RWR achieved the opposite effect than desired.

We then applied the three TAD detection algorithms, CaTCH, HiCseg, and TopDom, to ascertain whether there was an improvement on their performance after the data matrix was smoothed (Figures 1, S2, S3; parameter settings for running these algorithms were given in Supplementary Table S2). For all data matrices considered, smoothed or not (and even for the underlying idealized matrix *P* without noise), CaTCH simply called every two consecutive loci a domain, leading to a total of 100 domains (tiny green boxes) and a corresponding ARI of 0.024. On the other hand, HiCseg was able to detect the last four, except the first, domains (blue boxes) for the ideal, observed and KR-normalized matrices.

Even though the domain structures are not visually apparent, HiCseg still managed to identify the last four domains for RWS2s and RWS3s. However, it started to falter when a larger number of steps was taken, as also characterized by the decreasing ARI values. For RWR, regardless of the restart probabilities, HiCseg erroneously called every two consecutive loci as a domain just like CaTCH, not surprisingly given prominent diagonal feature of the RWR-smoothed matrices. For TopDom, it correctly identified all the domains for the ideal, observed noisy, KR-normalized, and the RWS2s matrices (purples boxes). However, as the number of steps increases, the performance degraded quickly. On the other hand, TopDom was able to identify the domains correctly for all the restart probabilities investigated except when it was very small (*α* = 0.05, in which case it broke the largest domain into two smaller ones).

These results, from a contrived dataset with known underlying ground truth, show that neither RWS nor RWR achieved the desired effects: they in fact destroy the domain structures, consistent with our theoretical results. Compared across the three domain detection algorithms, TopDom is seen to be the most successful, although not perfect. Specifically, CaTCH failed completely even with the ideal data without any noise. The performance of HiCseg and TopDom appear to be complementary in some sense: while HiCseg has a bit more success with some RWS-smoothed data, TopDom achieves much better results with the RWR-smoothed data.

### 3.1.2 Study 2: Biophysical-law-based simulated Hi-C data

It is well-known that chromatin interaction frequencies are inversely related to genomic distance, referred to as the biophysical law, or the power-decay law [1, 35]. In Hi-C data analysis, the logarithm of an interaction frequency between two loci is therefore typically taken to be linearly related to the genomic distance between these two loci [36]. The simulated data in this study observe this biophysical law in the background count matrix as we described in the following. First, a matrix *A* of size *N* × *N* was created following the power-decay law: the (*i, j*)^th^ element *A*_*ij*_ represents the expected contact count between loci *i* and *j*: log(*A*_*ij*_)= *c* + *d* log(*j* − *i*), 1 ≤ *i* < *j* ≤ *N*, where *c* and *d* are constants, with *d* being a negative number to observe the power-decay law and *c* a positive number to control the sequencing depth [37]. For *i* = *j*, that is, the counts on the diagonal, we set log(*A*_*ij*_)= *c*_1_, with *c*_1_ set to be at least as large as *c*. In our simulation, we set *c* = *c*_1_ = log(20), *d* = −0.4, and *N* = 1000. Then, the background matrix *B* was generated by sampling from a negative binomial (NegBin) distribution: *B*_*ij*_ ~ NegBin (mean = *A*_*ij*_, variance = *z* × *A*_*ij*_), 1 ≤ *i* ≤ *j* ≤ *N*, and *B*_*ji*_ = *B*_*ij*_, for *i* > *j*, producing over-dispersed data when *z* > 1. In our study, we set *z* equal to 1.2. This background matrix can be interpreted as the expected count matrix generated according to the biophysical law plus noise to induce features such as overdispersion. We then added a domain structure by creating a count matrix *D*: we first randomly divided the *N* = 1000 loci into *k* blocks, with each block serving as a domain and denoting the size as *n*_1_, …, *n*_*k*_. Then for each element (*i, j*) in each diagonal block submatrix of size *n*_*b*_ × *n*_*b*_, *b* = 1, …, *k*, we assigned it a domain effect from a Poisson distribution: *D*_*ij*_~ Poisson(*λ*_*b*_*B*_*ij*_), where *λ*_*b*_ was generated from Unif(*l, u*). In our simulation, we set *k* = 20, *l* = 0.2 and *u* = 0.7. Note that *λ*_*b*_ is block, not position, dependent; therefore, the *D* matrix is symmetric. All elements in the rest of the matrix (i.e. not in any of the *k* diagonal block submatrices) are set to be 0. Combining information from the background and the domain matrices, we have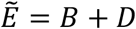. Finally, we account for Hi-C matrix sparsity by randomly setting *π*_1_ × 100*%* of the within-domain elements and *π*_2_ × 100*%* of those outside of any domain in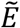to be zero, leading to our observed contact matrix *E*. In our simulation, we let *π*_1_ = 0.1 and *π*_2_ = 0.2.

The heatmaps for one simulated count matrix, *E*, and its KR-normalized counterpart *E*_*KR*_, are provided in Supplementary Figures S4 and S5, respectively, where the true domains (red boxes) are shown by superimposing them on the data matrices. Compared to the counterparts for the first study, the TADs are less obvious with the raw data; thus, a harder problem. After applying RWS with two steps, the domains are still somewhat visible, but almost completely invisible with a larger number of steps. For RWR with any restart probabilities, the most prominent feature of the matrices is the diagonal with much higher contact frequencies than the rest.

We once again apply the three domain detection algorithms to discern whether RWS or RWR smoothing can lead to better TAD identification (Figures S4 and S5). As with the first example, CaTCH failed to correctly identify any domains. With the raw count data matrix *E*, CaTCH did not identify any TAD (i.e. all loci are lumped into one group), resulting in an ARI of 0. For the KR-normalized data matrix and all the RWS- and RWR-smoothed matrices, CaTCH simply called every consecutive pair of loci as a TAD — like in the first example — thus failed to detect any of the 20 underlying TADs; this is reflected in a very low ARI of 0.026. For HiCseg, with the observed count data *E*, many of the underlying TADs were correctly identified, leading to a high ARI of 0.90. However, with the KR-normalized matrix, it only achieved an ARI of 0.53, since many of the larger true TADs were broken into smaller ones. For the RWS-smoothed matrices, some of the TADs were recovered; however, the performance degrades with more steps. The 3-step result, as recommended in GenomeDISCO [16], only achieved an ARI of 0.47. While the ARI for the RWS2s matrix is improved to 0.63, it is still much worse than the observed matrix without normalization nor smoothing. Regardless of the restart probability, HiCseg was unable to recover any meaningful (underlying) TADs, leading to the ARI all about 0, like CaTCH. TopDom for the observed and KR-normalized matrices led to a respectable ARI of 0.61 and 0.63, respectively, recovering many of the true TADs. Its performance improved to 0.76 for RWS3s. With taking more steps of the random walk, however, the results degraded like HiCseg, and TopDom was unable to identify meaningful domains with five or more steps. For RWR-smoothed matrices, the performance of TopDom is incredibly stable across the different restart probabilities, with almost identical results and an ARI of 0.75 for relatively small restart probabilities (*α* = 0.05, 0.1, 0.2).

We carried out the above simulation 100 times to gauge variability across multiple datasets (Figure 2). We see that CaTCH has no ability for detecting the underlying TADs, consistent with all results that we have seen thus far. For HiCseg, the original count matrix in fact has the best results. For the RWS-smoothed matrices, the performance was the best for taking a two-step random walk — which was generally better than the KR-normalized matrix without smoothing — but degraded as the number of steps increases. For the RWR-smoothed matrices, HiCseg’s performance was lacking regardless of the restart probability. From these results, it is observed that, while RWR-smoothed matrices do not lead to a greater power for detecting the underlying TADs, RWS-improved matrices can lead to better TAD detection power, but the number of steps is critical. Using TopDom, we see that the original count data, the KR-normalized matrix, the RWS-smoothed matrices with two or three steps, and all RWR-smoothed matrices regardless of the restart probabilities, all led to similarly good performances.

**Figure 2.**
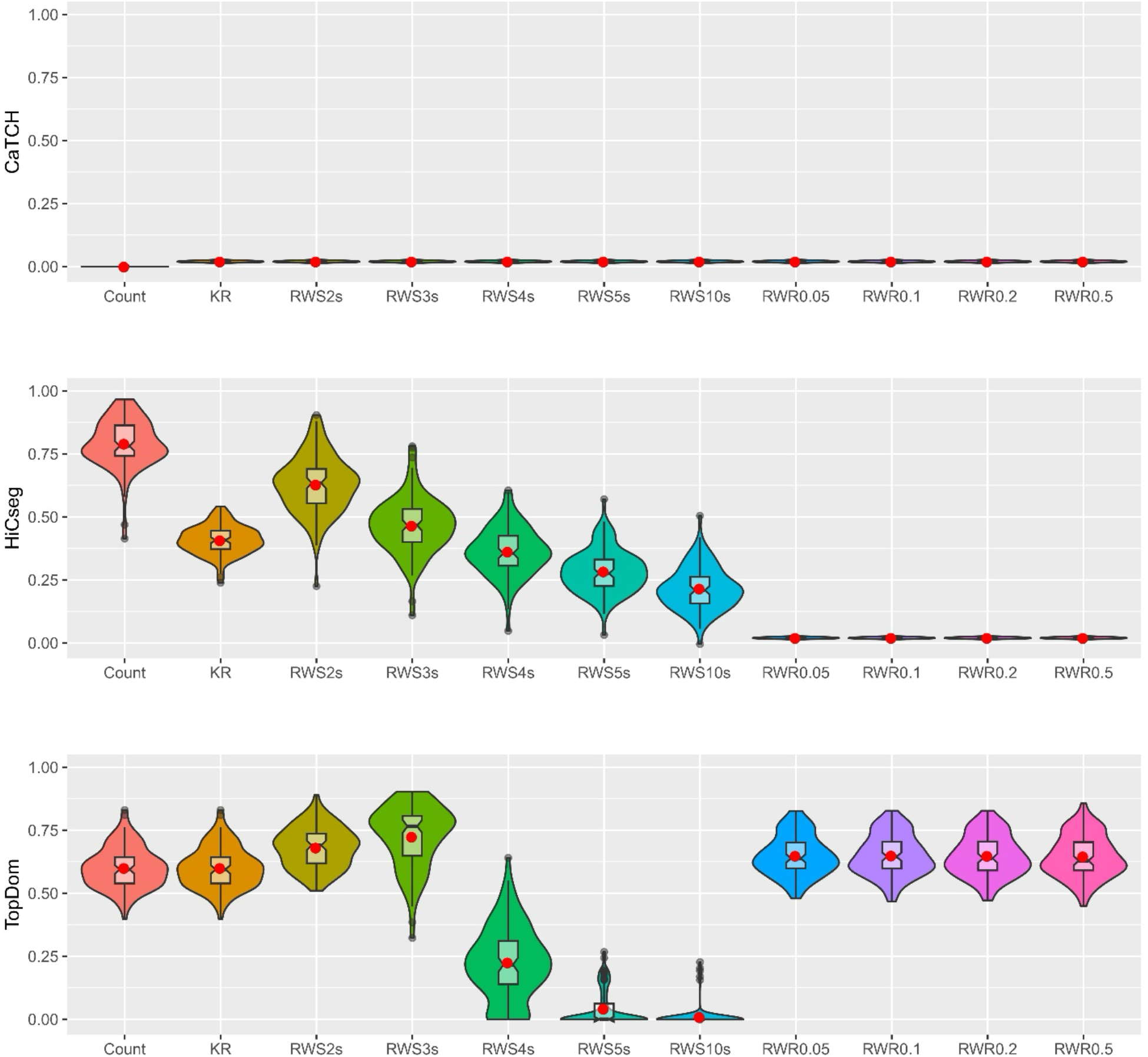
Violin plots of the ARI values, based on 100 replications, from the results of three TAD detection algorithms. Eleven kinds of data matrices were considered: count data, KR-normalized data, five RWS-smoothed matrices, and four RWR-smoothed matrices. Results are shown for CaTCH (row 1), HiCseg (row 2), and TopDom (row 3).

These results indicate that, overall, RWR-smoothed matrices can lead to slightly better performance for the identification of TAD boundaries if an appropriate algorithm is selected. For RWS-smoothed matrices, there is a potential for better detection performance, although the result is highly dependent on the selection of the number of random walk steps as well as the TAD detection algorithms. Finally, we observed that, although there is variability, the result seen for a single dataset appears to carry over from replicate to replicate.

#### 3.1.3 Study 3: Subsampling from a bulk Hi-C dataset

Although the first two studies provide a good avenue for evaluating the RWS- and RWR-smoothed matrices and TAD detection algorithms since the underlying ground truth is known, they are nonetheless not based on real data, even though the data in the second study were simulated observing the biophysical law. Therefore, in this study, we subsampled from a real bulk dataset to obtain contact matrices that are more in line with real single cell data to evaluate the random-walk algorithms and TAD detection methods for single cells, where the sequencing depth is usually only up to one-tenth of that for bulk data. Specifically, we considered the long arm of chromosome 22 of the K562 bulk A dataset with 200 kb resolution (https://www.ncbi.nlm.nih.gov/geo/query/acc.cgi?acc=GSM2109887). We subsampled 10% of the bulk data by using a multinomial distribution, where the probability of obtaining a contact at each location of the matrix (in the upper triangular) is proportional to its original observed count. The counts for the lower triangular matrix are set by flipping the upper triangular matrix to obtain symmetry. Since RWS and RWR were originally used to address the problem of sparsity in single cells, we are particularly interested in assessing the performance in this subsampling setting by treating each subsampled as a “single cell.” Although there is no ground truth in terms of TADs, one may compare the performance of TAD detection between the bulk data and the subsampled data to see whether there is any improvement in congruence with the bulk data for RWS- and RWR-smoothed matrices.

Using the count data directly for one sample, we can see that there are some consistencies in domain structures between the bulk and the subsampled data detected by HiCseg (ARI = 0.62) and CaTCH (ARI = 0.48) (Figure S6). However, TopDom lumped all loci after the first few together into a single domain, leading to an ARI of almost 0. We then KR-normalized both the bulk and the subsampled data. It is interesting to see that KR-normalized subsampled data has even less resemblance with the bulk data in terms of domain structures (Figure 3(a)), although we see that the TADs detected by TopDom for the KR-normalized bulk data and the raw bulk data before normalization are almost identical, which is not true for the other two domain detection algorithms. We then further obtained RWS- and RWR-smoothed data based on the KR-normalized subsample matrix. We see that some domain structures become visible with the RWS-smoothed matrices, especially with a larger number of steps (Figures 3(a) and S7). RWR with a small restart probability, 0.05, 0.1, and 0.2, also seemed to reveal some domain structures; however, the diagonal feature started to take over with a larger restart probability (Figure S8). Since the underlying ground truth is unknown, one could not tell for sure whether the domain structures visually apparent were true TADs, especially given that such structures were not visible in the bulk data. Nevertheless, it is interesting to see that, although TopDom was not able to find any TADs with the subsampled data, it detected TADs for all RWS- and RWR-smoothed matrices, and all led to a higher ARI with the bulk data. For HiCseg, it called many meaningless small domains for all the data matrices (including the bulk data) except for RWS10s. Not surprisingly, CaTCH failed again, calling every two consecutive bins as a domain, for all the normalized and smoothed matrices.

**Figure 3.**
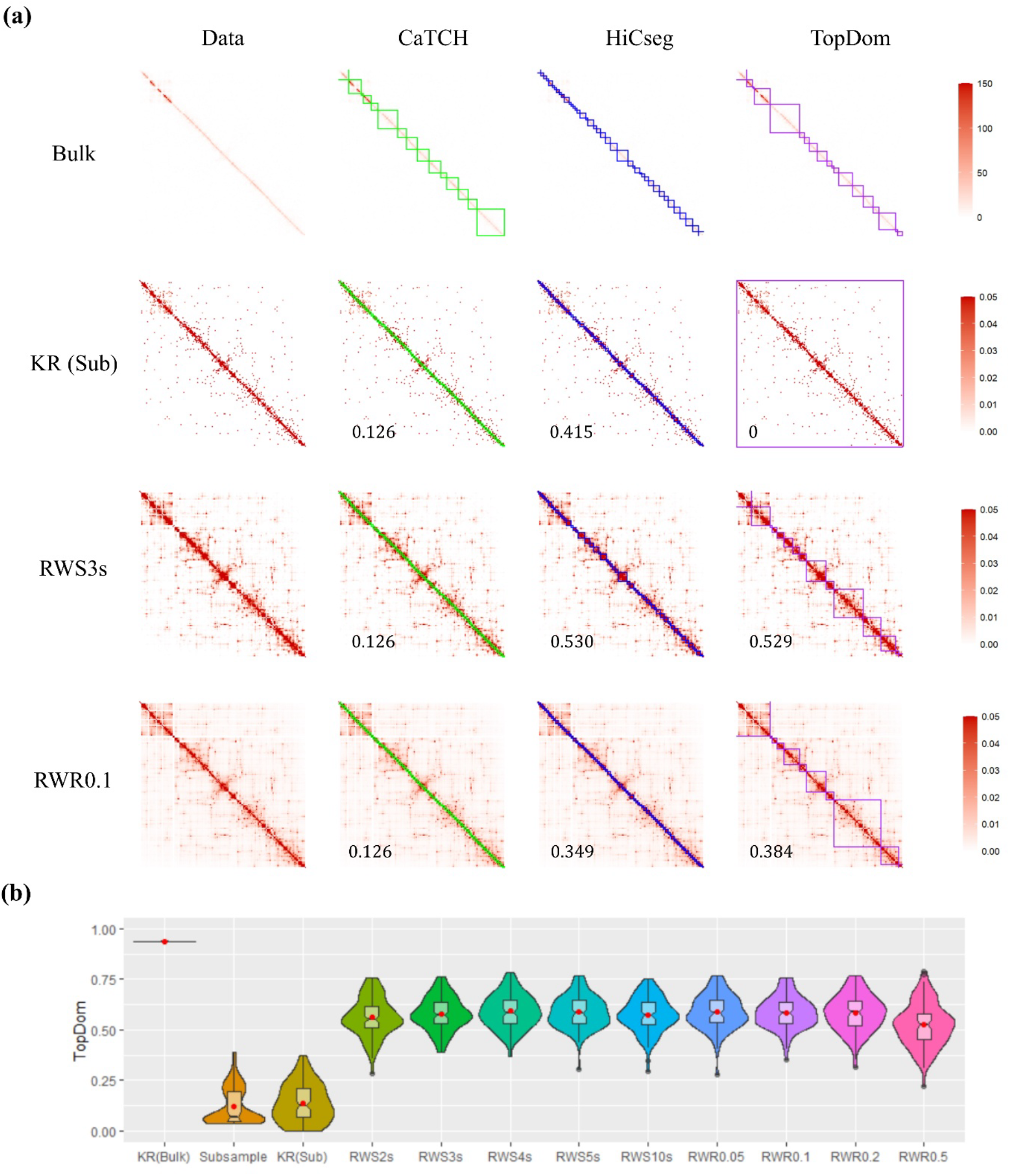
Results based on the subsampling procedure to simulate single cell data. (a) Data matrices (column one) and detected domain boundaries (columns 2-4 for CaTCH, HiCseg, and TopDom, respectively); row 1 shows the original bulk count matrix, while rows 2-4 provide, respectively, the KR-normalized, SWS3s, RWR0.1, results from one subsample. Note that 150 and 0.05 are the censured values for the heatmaps. (b) Violin plots of ARI values for the results from TopDom with 11 kinds of data matrices with 100 replicated subsamples.

To gauge variability, we performed the subsampling 100 times, and we plotted the distributions of the ARIs between the subsampled matrices (before and after various levels of smoothing) and the bulk data. The results for TopDom applying to the RWS- and RWR-smoothed data are fairly consistent across a wide range of number of steps and restart probabilities, and there is an appreciable improvement over the results for the subsampled count or KR-normalized subsampled data matrices (Figure 3(b)). However, for CaTCH and HiCseg, the results from the count matrix are better than their KR-normalized and smoothed counterparts (Figure S6(b)).

### 3.2 Analysis of two experimental Hi-C datasets

We now turn to experimental data to further substantiate our findings from the three simulation studies. We evaluated whether the RWS- and RWR-smoothed matrices from real experimental data can lead to improvements on TAD detection using the same three algorithms that we investigated in the simulation studies. To be comprehensive, we considered two publicly available Hi-C datasets — one bulk and one single cells — that provides a range of data sparsity.

#### 3.2.1 Human Embryonic Stem Cells Bulk Hi-C Data

We considered a bulk Hi-C dataset downloaded from the public domain (https://www.ncbi.nlm.nih.gov/geo/query/acc.cgi?acc=GSE35156). This human Embryonic Stem Cells (hESC) Hi-C dataset was originally generated for defining and studying TADs [4]. We used the 40 kb resolution processed data [30] and focused on the 857 loci in Chromosome 22. Although there are a total of 1243 loci in the chromosome, the contact counts among the first 386 (the short arm and the centromere) are all zeros; therefore, they were not included in our analysis. The original data matrix exhibits visible domains, but such information becomes blurrier as the number of steps increases in RWS (Figure S9). With different restart probability, contact information other than the main diagonal also get diluted (Figure S10). Interestingly, CaTCH was able to detect domains that are obviously visible for the unprocessed data, but failed to provide any meaningful domain information for the rest of the data matrices. HiCseg also detected many domains, but at a finer scale for some than those called by CaTCH. HiCseg continues to detect domains for the RWS-smoothed matrices, but at a coarser and coarser scale as the number of steps increases. As we have seen with the simulated and subsampled data, HiCseg failed to detect anything meaningful with any of the RWR-smoothed matrices. For TopDom, the domain boundaries detected based on the raw data is remarkably similar to those from CaTCH. For the RWS-smoothed matrices, its behavior tracked those produced by HiCseg: as the number of steps increases, the domains detected becomes coarser and coarser, and eventually becoming meaningless with RWS10s. However, unlike the other two TAD detection algorithms, TopDom detected some likely-meaningful domains with the RWR-smoothed data matrices, although the inherent diagonal feature of the smoothed matrices led to many small domains.

Overall, we observe some consistency among domain structures inferred by the same TAD calling algorithm and among the same type of smoothing methods. Further, the domain structures inferred by HiCseg based on the RWS-smoothed matrices, apart from RWS10s, are similar to those detected by TopDom, regardless of whether they were inferred from RWS- or RWR-smoothed matrices (Figure 4). However, the domain structures detected by CaTCH (or the lack of) do not share much commonality with those from HiCseg or TopDom.

**Figure 4.**
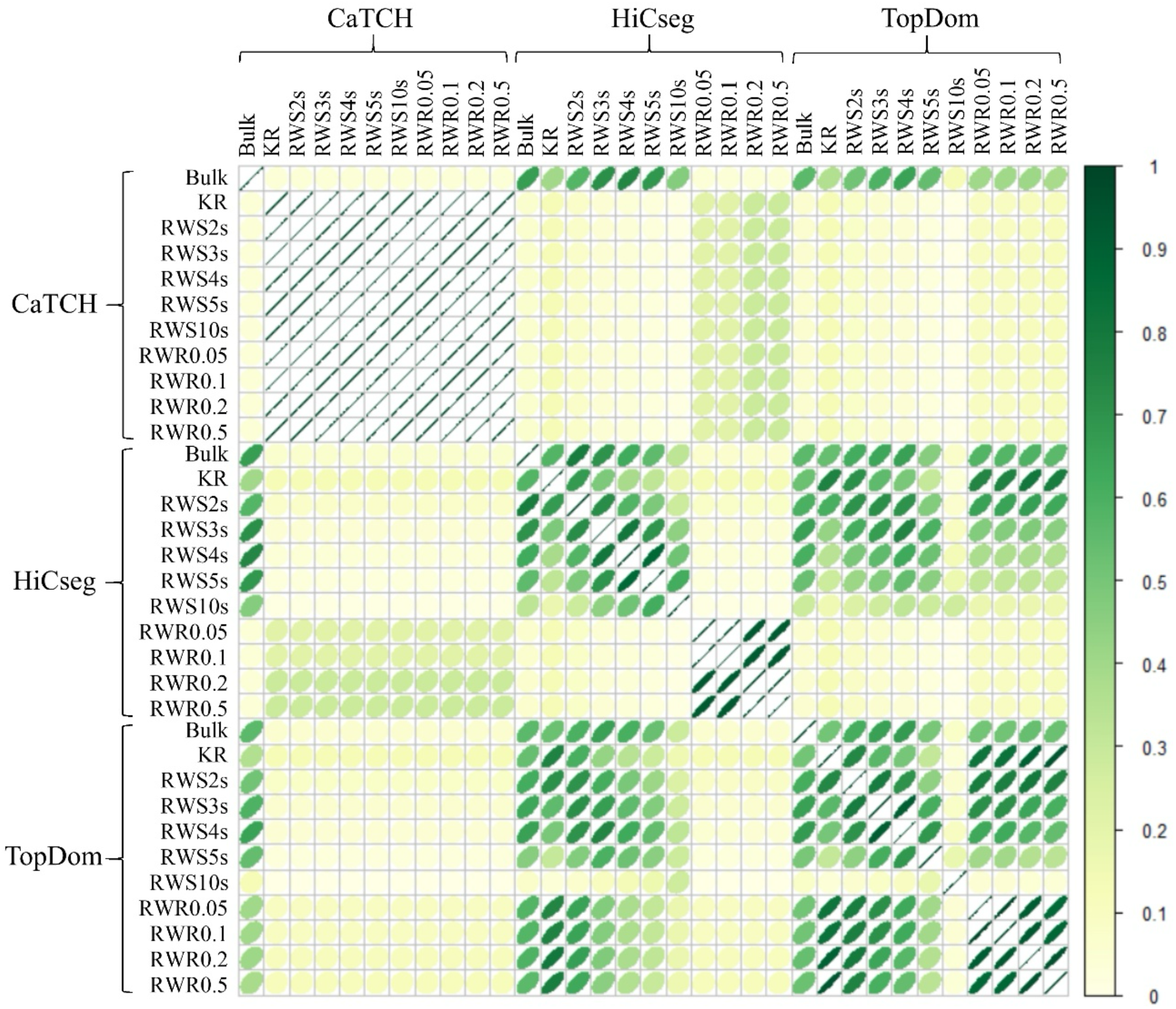
Correlation plot of the ARI values between the identified domain boundaries on the bulk count, KR-normalized, and RWS-/RWR-smoothed matrices of the hESC data. We considered 11 kinds of matrices (“bulk”, KR-normalized, five RWS-smoothed, and four RWR-smoothed matrices) and three TAD detection methods (CaTCH, HiCseg, and TopDom).

#### 3.2.2 GM and PBMC Single Cell Hi-C Data

In our second real data analysis, we considered a single-cell Hi-C dataset (https://www.ncbi.nlm.nih.gov/geo/query/acc.cgi?acc=GSE117874) consisting of 14 lymphoblastoid (GM) and 18 peripheral blood mononuclear cells (PBMC) samples. We used chromosome 22 contact counts at 40 kb resolution, leading to a total of 860 loci after excluding those that have interaction counts being 0 (short arm and centromere). For evaluating the performance, we created a composite contact matrix combining the counts from all single cells for each of the two types, assuming intra-type similarity. The domains detected from the composite contact matrices will be treated as the “gold standard” for comparing with those from each of the single cells within the same type. For the GM cells, CaTCH’s analysis on the original single cells counts matrices led to reasonable congruence with the gold standard, with most of them achieving an ARI of more than 0.6. However, none of the smoothed matrices produced sensible results, with ARI being close to 0 (Figure 5(a)). For HiCseg, the best result of congruence is also achieved with the original single-cell count matrix, although RWS with an appropriate number of steps (RWS5s or RWS10s) also led to similarly reasonable performance. For TopDom, there is a large discrepancy between the “gold standard” and the domains obtained from the original single-cell count matrix, which might be due to the detection of some large domains in single cells (Fig S11). In contrast, the RWS- and RWR-smoothed matrices all led to reasonably good results. Overall, these results are consistent with what we observed in our simulation studies. For the PBMC cells, the results are qualitatively the same as those for the GM cells (Figures 5(b) and S11).

**Figure 5.**
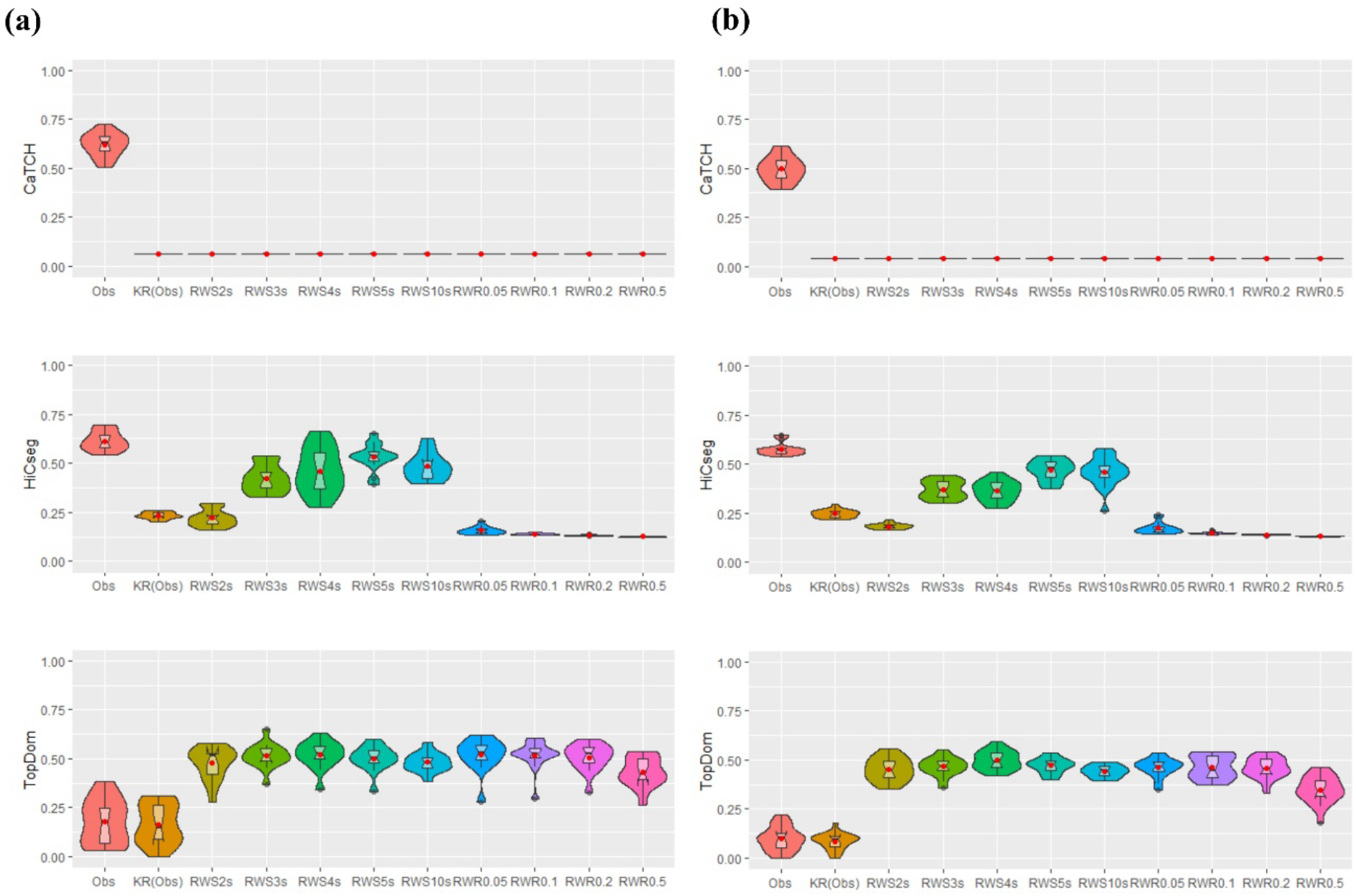
Violin plots for the ARI values of the results from CaTCH (row 1), HiCseg (row 2), and TopDom (row 3). (a) 14 GM cells; (b) 18 PBMC cells. Each ARI value was computed by comparing the domain boundaries detected by the composite count matrix (combining all data) with each of the single cells.

## 4 Discussion and Conclusion

In this paper, we studied whether random-walk-based methods, RWS and RWR, are appropriate tools for improving single-cell Hi-C data quality. These methods are often used as part of a pipeline for Hi-C data analysis, especially for single-cell Hi-C data, yet little justification is available in the literature. Our extensive investigation, through theoretical analyses, simulation studies, and real data applications all converge to the following findings: RWS-smoothed data matrices will lead to a “monolith” (all entries have the same contact probabilities) if too many steps are taken; RWR-smoothed data, regardless of its restart probability, will lead to the prominent diagonal feature. Therefore, in both types of random-walk-based smoothing approaches, the smoothed matrix may lose the underlying TAD structures rather than enhancing them. In almost all the simulation and real data studies, using the original count matrix usually led to comparable, if not better, performance with the best smoothed data, with a few exceptions.

As a side product, our study also provided an evaluation of three TAD detection algorithms, which were deemed as the top performers in a previous study. Among the three TAD detection algorithms, CaTCH was seen to be least competitive, failing to uncover the underlying TADs most of the time, even under the ideal TAD structure setting regardless of whether the original count or normalized/smoothed matrices are used. HiCseg often has good performance with the original count data, and using RWS-smoothed data with an appropriate number of steps can sometimes lead to improved TAD detection capability. However, HiCseg typically failed to detect TADs with RWR-smoothed matrices. TopDom was seen to have the best overall performance. It can often produce results that are at least comparable, and sometimes better than using the original count matrix, if an appropriate number of steps for RWS or an appropriate restart probability for RWR were used.

In conclusion, the results from our extensive study do not support the use of random walk as a data improvement technique. Nevertheless, if a reader would still choose to use a random-walk based method, then an important issue that deserves further discussion is the need for doubly stochastic matrix before applying RWS and RWR. Although a transition probability matrix is sufficient for a random walk, one without symmetry destroys the inherent feature of a contact matrix. Of the commonly used normalization methods, including ICE, symmetry is not guaranteed after normalization. In contrast, KR normalization has the capability of turning a symmetric matrix into a doubly stochastic matrix, in which not only each row sums to 1, but each column sums to 1 also.

## Acknowledgement

This work was supported in part by the National Institutes of Health [R01GM114142 to SL].

## Notes

### Competing Interest Statement

The authors have declared no competing interest.

